# CosMIC: A Consistent Metric for Spike Inference from Calcium Imaging

**DOI:** 10.1101/238592

**Authors:** Stephanie Reynolds, Therese Abrahamsson, P. Jesper Sjöström, Simon R. Schultz, Pier Luigi Dragotti

## Abstract

In recent years, the development of algorithms to detect neuronal spiking activity from two-photon calcium imaging data has received much attention. Meanwhile, few researchers have examined the metrics used to assess the similarity of detected spike trains with the ground truth. We highlight the limitations of the two most commonly used metrics, the spike train correlation and success rate, and propose an alternative, which we refer to as CosMIC. Rather than operating on the true and estimated spike trains directly, the proposed metric assesses the similarity of the pulse trains obtained from convolution of the spike trains with a smoothing pulse. The pulse width, which is derived from the statistics of the imaging data, reflects the temporal tolerance of the metric. The final metric score is the size of the commonalities of the pulse trains as a fraction of their average size. Viewed through the lens of set theory, CosMIC resembles a continuous Sørensen-Dice coefficient — an index commonly used to assess the similarity of discrete, presence/absence data. We demonstrate the ability of the proposed metric to discriminate the precision and recall of spike train estimates. Unlike the spike train correlation, which appears to reward overestimation, the proposed metric score is maximised when the correct number of spikes have been detected. Furthermore, we show that CosMIC is more sensitive to the temporal precision of estimates than the success rate.

## 1 Introduction

Two-photon calcium imaging has enabled neuronal population activity to be monitored in vivo in behaving animals (Dombeck et al., 2010; Peron et al., 2015). Modern microscope design allows neurons to be imaged at sub-cellular resolution in volumes spanning multiple brain areas (Sofroniew et al., 2016). Coupled with the current generation of fluorescent indicators (Chen et al., 2013), which have sufficient sensitivity to read out single spikes, this imaging technology has great potential to further our understanding of information processing in the brain.

The fluorescent probe, however, does not directly report spiking activity. Rather, it reads out a relatively reliable indicator of spiking activity — a cell s intracellular calcium concentration — from which spike times must be inferred. A diverse array of techniques have been proposed for this task, including deconvolution approaches (Vogelstein et al., 2010; Friedrich et al., 2017; Pachitariu et al., 2017), methods that identify the most likely spike train given a signal model (Vogelstein et al., 2009; Deneux et al., 2016) and approaches that exploit the sparsity of the underlying spike train (Oñativia et al., 2013). To enable the investigation of neural coding hypotheses, reconstructed spike trains must have sufficient temporal precision for analysis of synchrony between neurons and behavioural variables (Huber et al., 2012), whilst accurately inferring the rate of spiking activity.

Although the development of spike detection algorithms has received a lot of recent attention, few researchers have examined the metrics used to assess an algorithm s performance. At present, there is no consensus in the best choice of metric. In fact, from our survey, 44% of papers presenting a new method assess its performance using a metric unique to that paper. This inconsistency impedes progress in the field — algorithms are not directly comparable and, consequently, data collectors cannot easily select the optimal algorithm for a new dataset.

The two most commonly used metrics, the spike train correlation (STC) and the success rate, are not well-suited to the task. The STC, which is invariant under linear transformations of the inputs, is not able to discriminate the similarity of the rates of two spike trains (Paiva et al., 2010). Moreover, the temporal binning that occurs prior to spike train comparison impairs the STC s ability to compare spike train synchrony (Paiva et al., 2010). These limitations suggest that the STC, whilst a quick and intuitive method, is not appropriate for assessing an algorithm s spike detection performance. The success rate, which does accurately compare spike rates, does not reward increasing temporal precision above a given threshold. Consequently, it is not an appropriate metric for evaluating an algorithm s performance when the end goal is, for example, to investigate the synchrony of activations within a network.

In this paper, we present a metric that can discriminate both the temporal and rate precision of an estimated spike train with respect to the ground truth spike train. Unlike the STC, we do not bin the spike trains. Rather, spike trains are convolved with a smoothing pulse that allows comparison of spike timing with an implicit tolerance. The similarity between the resulting pulse trains is subsequently assessed. This type of continuous approach is also preferred by metrics assessing the relationship between spike trains from different neurons (van Rossum, 2001; Schreiber et al., 2003). We set the pulse width to reflect the temporal precision that an estimate is able to achieve given the statistics of the dataset. As such, the metric is straightforward to implement since there are no parameters to tune. For convenience, we refer to the proposed metric as CosMIC — a Consistent Metric for spike Inference from Calcium imaging. In the following, we demonstrate CosMIC s ability to discriminate spike train similarity on real and simulated data. We include comparisons against the two most commonly used metrics, the spike train correlation and the success rate, and against two metrics designed to assess similarity between spike trains from different neurons (Victor and Purpura, 1997; van Rossum, 2001).

## 2 Constructing the metric

In this paper, we present a metric for comparing the similarity of two sets of spikes: a ground truth set, 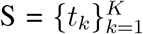, and a set of estimates, 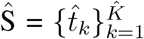. Due to limiting factors, such as noise and model mismatch, it is improbable that an estimate will match a true spike with infinite temporal precision. As such, we do not expect that *t̂*_*j*_ = *t*_*k*_ for any *j* or *k*. Rather, we wish to reward estimates within a reasonable range of accuracy given the limitations of the data. We achieve this by leveraging results from fuzzy set theory (Zimmermann, 2010).

In contrast to classical sets, to which an element either belongs or does not belong, fuzzy sets contain elements with a level of certainty represented by a membership function — the higher the value of the membership function, the more certain the membership. In the following, we define two fuzzy sets, Ŝ_*ϵ*_ and Ŝ_*ϵ*_, which represent the original sets of spikes, S and Ŝ, with a level of temporal tolerance defined by a parameter *ϵ*. We set *ϵ* to reflect the temporal precision that an estimate is able to achieve given the statistics of a dataset (see Section 3). The corresponding membership functions *y*(*t*) and *ŷ*(*t*), which are defined for *t* ∊ ℝ, are calculated through convolution of the spike trains,

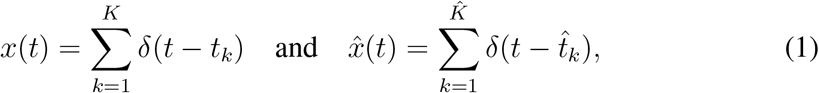

with a triangular pulse, *p*_*ϵ*_(*t*), such that *y*(*t*) = *x*(*t*) ∗ *p*_*ϵ*_(*t*) and *ŷ*(*t*) = *x̂*(*t*) ∗ *p*_*ϵ*_(*t*). The resulting functions have local maxima at the locations of the respective sets of spikes (Fig. 1A). As *x*(*t*) and *x̂*(*t*) are analogous to the membership functions of the classical sets of spikes, we can think of the convolution as a temporal smoothing of the membership. The pulse that we employ is a triangular B-spline (Fig. 1B),

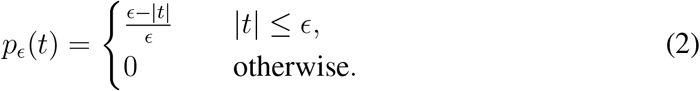

Using this triangular pulse means that, the further a time point, *t*, is from a spike, the less weight the membership function receives at that point. Past a certain distance, *ϵ*, the membership function receives no weight. Many pulse shapes could be chosen to introduce this grading of temporal precision, we select a triangular pulse as it is straightforward to examine analytically and implement computationally.

**Figure 1:**
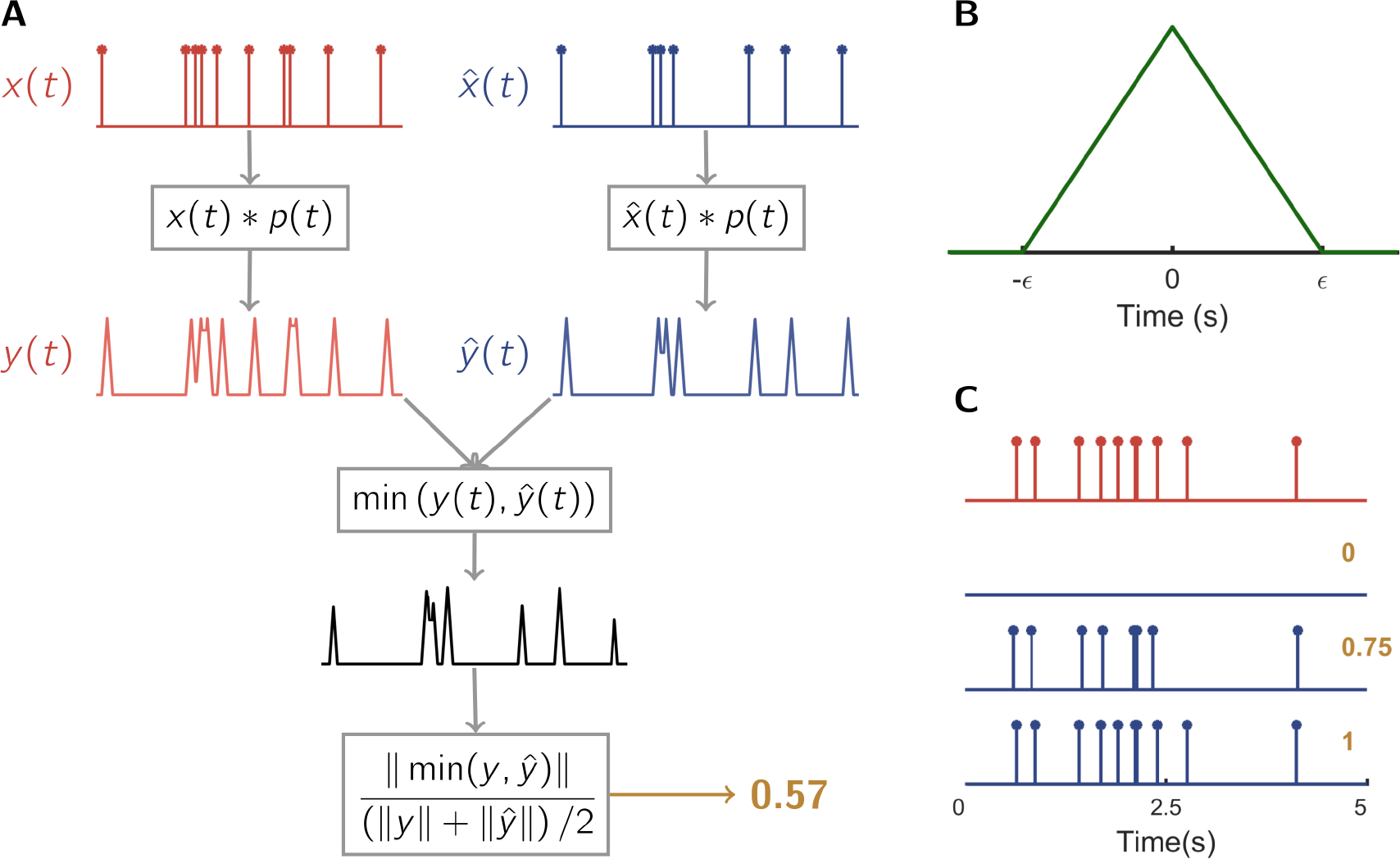
A flow diagram of the proposed metric. The ground truth spike train and estimated spike train are convolved with a triangular pulse (**B**), whose width is determined by the statistics of the data. The metric compares the difference between the resulting pulse trains (**A**). Metric scores are in the range [0,1] — a perfect estimate achieves score 1 and an empty spike train is scored 0 (**C**).

We design the proposed metric to quantify the size of the intersection of the fuzzy sets of true and estimated spikes with respect to the average size of the sets, such that

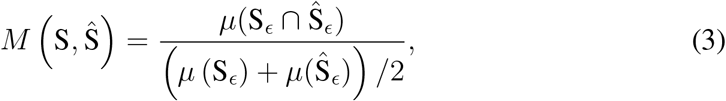

where *μ* is the L1-norm: *μ*(S_*ϵ*_) = ‖*y*‖ = ∫_ℝ_|*y*(*t*)| d*t*. An analogous formula was presented for discrete fuzzy sets by Pappis and Karacapilidis (1993). Our formula can be interpreted as the continuous version of the Sørensen-Dice coefficient (Dice, 1945; Sørensen, 1948) — a score which is commonly used to assess the similarity of discrete, presence/absence data. Also known as the F1-score, in the context of spike detection, the Sørensen-Dice coefficient is referred to as the success rate (Section 4.1).

The membership function of an intersection of sets is the minimum of their respective membership functions. It follows that

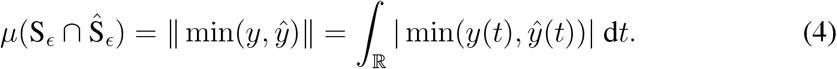

Taking the minimum of the membership functions produces a conservative representation of the intersection of two sets; in our context, spikes that appear in one spike train and not in the other are removed (Fig. 2A) and spikes that are detected with poor temporal precision are assigned less weight (Fig. 2B and 2C).

**Figure 2:**
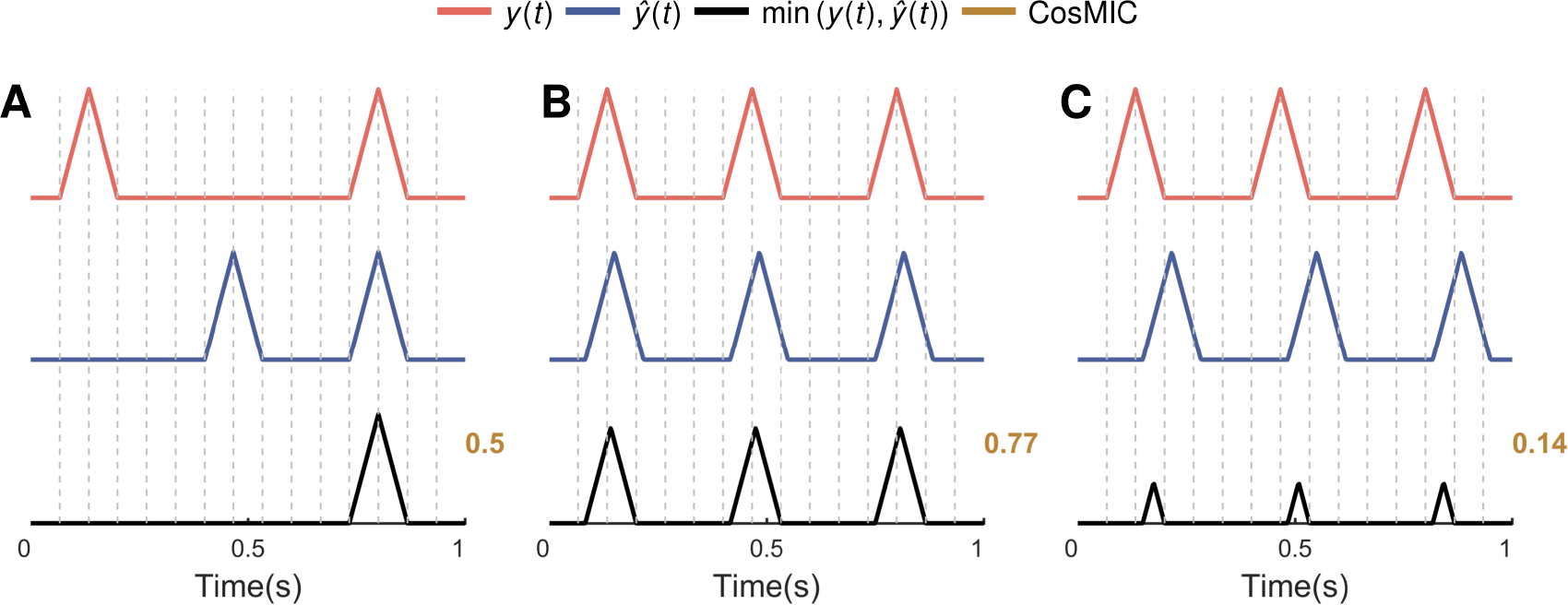
The proposed metric quantifies the commonalities of the sets of true and estimated spikes as a proportion of the average size of those sets. Commonalities are found by taking the minimum of the pulse trains — as such, spikes that appear in only one pulse train are excluded (**A**) and estimates with lower temporal precision receive a lower score (**B** and **C**).

The metric can also be written in alternative form

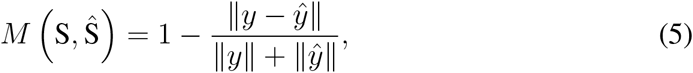

the derivation of which is shown in Appendix A.1. From Eq. (5), it is clear that the maximal score of 1 is achieved when the membership functions, and therefore the sets of true and estimated spikes, are equivalent. The minimal score of 0 is achieved when the support of the membership functions do not overlap, i.e. no estimates are within the tolerance of the metric (Fig. 1C).

### 2.1 Ancestor metrics

Like the success rate, CosMIC can alternatively be derived from a pair of metrics, which we refer to as ancestor metrics. The first of these metrics measures the proportion of ground truth spikes that were detected within the precision of the pulse width, such that

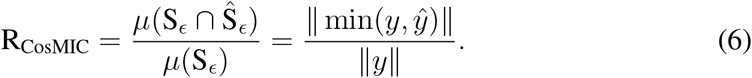

This score is analogous to the recall of a spike train estimate, one of the ancestor metrics from which the success rate is formed. The second of CosMIC’s ancestor metrics measures the proportion of estimated spikes that detect a ground truth spike within the precision of the pulse width, such that

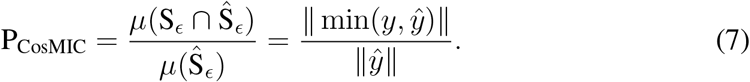

This is analogous to the precision, the second metric used to compute the success rate. Finally, computing the harmonic mean of the two ancestor metrics and rearranging, we obtain CosMIC:

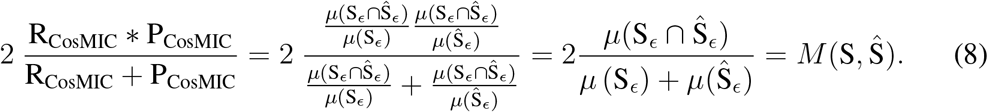

The analogy to the success rate can be seen clearly from the presentation of that metric in Section 4.1.

## 3 Temporal error tolerance

The width of the triangular pulse with which the spike trains are convolved reflects the accepted tolerance of an estimated spike s position with respect to the ground truth. To set this width, we calculate a lower bound on the temporal precision of the estimate of one spike — the Cramér-Rao bound (CRB) — from the statistics of the data. The CRB reports the lower bound on the mean square error of any unbiased estimator (Kay, 1993). It is therefore useful as a benchmark; an estimator that achieves the CRB should be awarded a relatively high metric score. In Section 3.1, we detail the calculation of the CRB. In Section 3.2, we outline how we use this bound to determine the pulse width. Then, in Section 3.3, we provide practical advice on the calculation of the bound.

### 3.1 Cramér-Rao bound for spike detection

We consider the problem of estimating the location of one spike, *t*_0_, from noisy calcium imaging data. The fluorescence signal is modelled as

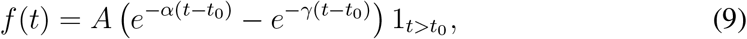

where *α*, *γ* and *A* are parameters that determine the shape and amplitude of the calcium transient. We assume that we have access to *N* noisy samples, such that

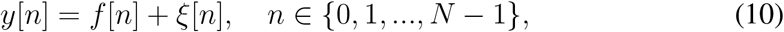

where *ξ*[*n*] are independent samples of a zero-mean Gaussian process with standard deviation *σ* and *f*[*n*] = *f*(*nT*) are samples of the fluorescence signal with time resolution *T*. The CRB on the uncertainty in the estimated position of *t*_0_ is

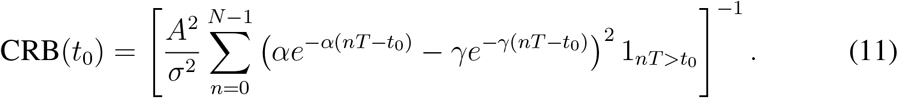

This bound was first presented by Schuck et al. (2017). The bound is derived by calculating the inverse of the Fisher Information, which, in the case of samples corrupted by independent, zero-mean, Gaussian noise, is

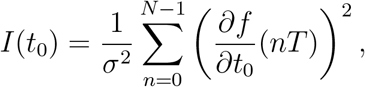

where ∂*f*/∂*t*_0_ is the derivative of the fluorescence signal with respect to the spike time, *t*_0_:

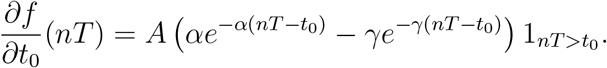

In this work, we use the CRB to set the temporal tolerance of the metric. In order that the CRB holds for an arbitrarily placed spike, we remove the dependency on the true spike time by averaging the result over several values of *t*_0_. We compute 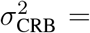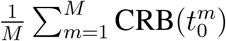, where 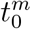 are evenly placed in the interval (*nT*, (*n*+1)*T*) for a fixed n. In Fig. 3, we plot *σ*_CRB_ as the sampling rate and peak signal-to-noise ratio (PSNR) of the data vary. The PSNR is computed as 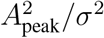, where *σ* is the standard deviation of the noise and *A*_peak_ is the peak amplitude (maximum) of the fluorescence signal in Eq. (9). For this example, we use *α* = 3.18s^−1^ and *γ* = 34.49s^−1^; the parameters for a Cal-520 AM pulse (Tada et al., 2014). We see that the CRB decreases as either the scan rate or the PSNR of the data increases.

**Figure 3:**
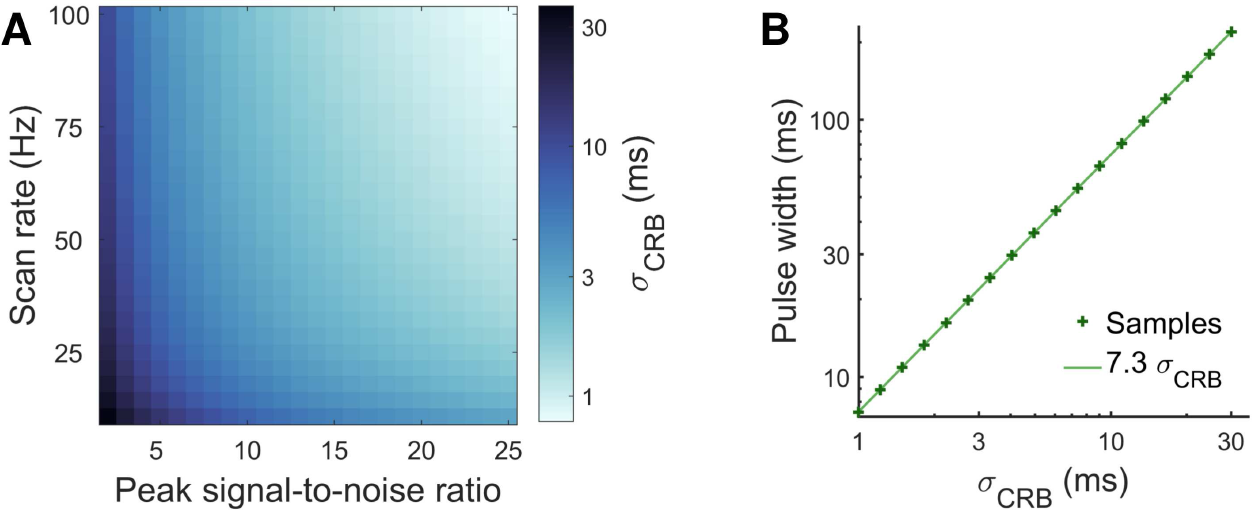
The pulse width is set to reflect the temporal precision achievable given the statistics of the dataset. We calculate the Cramér-Rao bound (CRB), 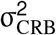, a lower bound on the mean square error of the estimated location of one spike from calcium imaging data (**A**). This bound decreases as the scan rate (Hz) and peak signal-to-noise ratio (squared calcium transient peak amplitude/noise variance) increase. We set the pulse width to ensure that an estimate of one spike at the temporal precision of the CRB achieves, on average, a score of 0.8. This results in a pulse width of approximately 7.3 σ_CRB_ (**B**).

### 3.2 Pulse width

The CRB can be used as a benchmark for temporal precision of any unbiased estimator. As such, we set the pulse width to ensure that, on average, an estimate at the precision of the CRB achieves a relatively high score. We set the benchmark metric score at 0.8, as this represents a relatively high value in the range of the metric, which is between 0 and 1. The importance of this score is not the particular benchmark value — there are a range of values that give similar performance — but rather that it is a reproducible number with a clear interpretation. In this paper, we characterise the discrimination performance of CosMIC with a benchmark value of 0.8, so that its scores can be interpreted when applied to spike inference algorithms on real data. The benchmark value was set lower than the metric s maximum value, 1, so that the score does not saturate when the model assumptions are not ideally satisfied. On real data, the noise may not be stationary (*σ* may vary in time), and so algorithms may appear to outperform the CRB. A benchmark score of 0.8 means that the metric score does not saturate in this scenario.

We consider a true spike at *t*_0_ and an estimate, *U*, normally distributed around it at the precision of the CRB, such that 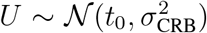. Then, we fix the pulse width so that, on average, 𝔼 [*M*(*t*_0_, *U*)] = 0.8. In Appendix A.3, we show that this condition is satisfied when

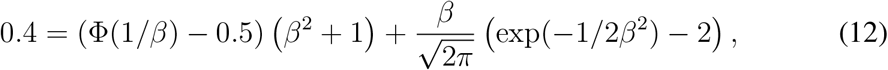

where *β* = *σ*_CRB_/*w*, *w* is the pulse width and Φ denotes the cumulative distribution function of the standard normal distribution. We observe that the pulse width that solves this equation is approximately equal to 7*σ*_CRB_ (Fig. 3B).

### 3.3 Implementation

Code to implement the metric can be found at github.com/stephanierey/metric along with a demonstration. In order to use the metric, one must have estimates of the fluorescence signal parameters, {*α*, *γ*, *A*, *σ*}, see Eq. (9). In the following, we provide some guidance on the estimation of these parameters. Alternative strategies have been suggested by numerous model-based algorithms, whose spike detection procedures utilise a subset of the above parameters (Vogelstein et al., 2009; Pnevmatikakis et al., 2013, 2016; Deneux et al., 2016).

The standard deviation of the noise, *σ*, can be computed as the sample standard deviation of a portion of the data in which there were no calcium transients. The parameters that determine the speed of the rise and decay of the pulse — *α* and *γ* — are predominantly defined by characteristics of the fluorescent indicator that was used to generate the imaging data. In Table 1, we provide documented values of *α* and *γ* for four commonly used fluorescent indicators, extracted from the corresponding references: Cal-520 AM (Tada et al., 2014), OGB-1 AM (Lütcke et al., 2013), GCaMP6f and GCaMP6s (Chen et al., 2013). These values can be used as a guideline; in practise, they will vary with the indicator expression level as well as the cell type. We note that the time taken for a calcium transient to rise to its peak and the decay time are functions of both *α* and *γ*; the values presented in Table 1 are thus not easily interpretable in terms of the shape of a calcium transient pulse.

**Table 1:** To calculate CosMIC s pulse width, the parameters that define the speed of rise and decay of the calcium transient, *α* and *γ*, are required. Here, we provide documented values of these parameters for four commonly used fluorescent indicators.

| Fluorescent indicator | $\alpha$ (s<sup>-1</sup>) | $\gamma$ (s<sup>-1</sup>) |
| --- | --- | --- |
| GCaMP6f | 4.88 | 60.97 |
| GCaMP6s | 1.26 | 15.16 |
| OGB-1 AM | 1.5 | 101.5 |
| Cal-520 AM | 3.18 | 34.39 |

It is typically necessary for a spike detection algorithm to estimate the value of the amplitude parameter, *A*, in order to detect spikes. Indeed, Vogelstein et al. (2009) integrate this step into the spike detection procedure, iteratively estimating the spike locations and the amplitude, amongst other parameters. If, however, *A* is not known, we recommend that the parameter is fit from the data samples and the signal model, such that

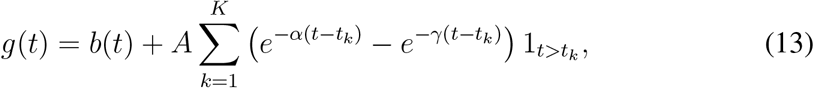

where *b*(*t*) is a baseline component and *α*, *γ* are the estimated pulse shape parameters. When the baseline component is constant and there is no indicator saturation, this is a linear problem. In practise, a neuron’s spike amplitude is not constant over time. In fact, depending on the fluorescent indicator, the amplitude may increase (Chen et al., 2013) or saturate (Lϋtcke et al., 2013) at high spike rates. We recommend that the amplitude parameter is fit from a subset of the data in which neither saturation nor supra-linear amplitudes are present.

## 4 Numerical experiments

To assess the discriminative ability of CosMIC, we simulate true and estimated spike trains in various informative scenarios. We compare CosMIC with the two most commonly used metrics in the spike inference literature, which we define in Sections 4.1 and 4.2 for completeness. We also compare against two metrics designed to assess the similarity of spike trains from different neurons. We define the metrics of Victor and Purpura (1997) and van Rossum (2001) in Sections 4.3 and 4.4, respectively.

### 4.1 Success rate

The success rate, which is defined as a function of the true and false positive rates or, alternatively, as a function of precision and recall, appears in various forms in the literature. Spike inference performance has been assessed using true and false positive rates (Rahmati et al., 2016), precision and recall analysis (Reynolds et al., 2017) and using the complement of the success rate, the error rate (Deneux et al., 2016). We study this class of metrics under the umbrella of the success rate, which we define here.

A ground truth spike is deemed to have been detected’ if there is an estimate within *δ*_1_/2(s) of that spike, where *δ*_1_ is a free parameter. Only one estimate can be deemed to detect one ground truth spike. The recall is the percentage of ground truth spikes that were detected. The precision is the percentage of estimates that detect a ground truth spike. Then, the success rate is the harmonic mean of the precision and recall, such that

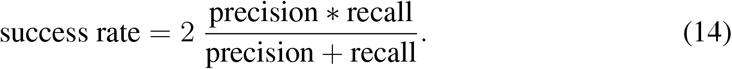

A binary true detection region centred around each ground truth spike is analogous to an implementation of CosMIC with a box function pulse. To ensure that the success rate ‘pulse’ has the same width as CosMIC s pulse, we set *Δ*_1_ = *ϵ*, where *ϵ* is half the pulse width, see Fig. 4.

**Figure 4:**
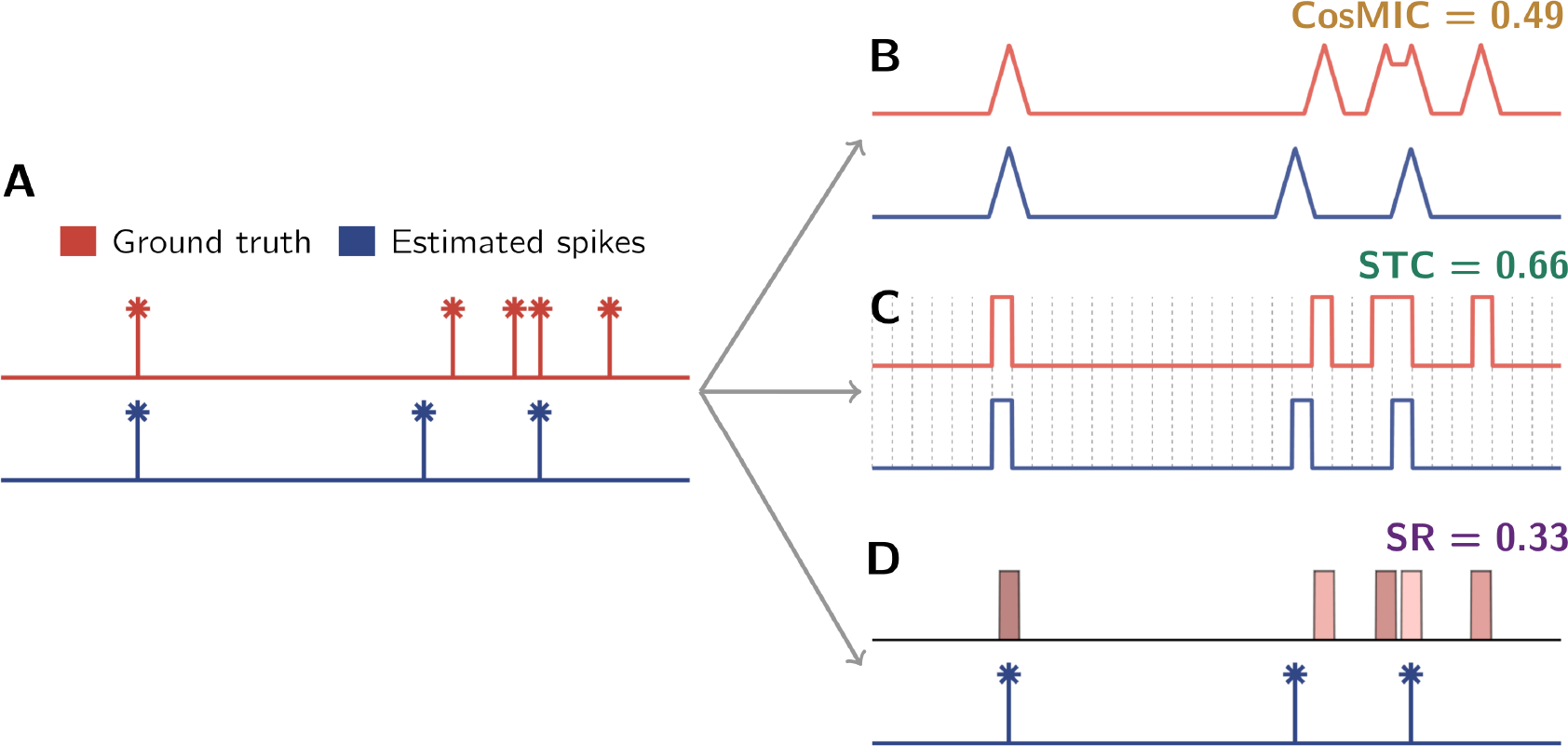
We compare the scores of three metrics: CosMIC, the spike train correlation (STC) and the success rate (SR). None of the metrics compute scores directly from the true and estimated spike trains, shown in **A**. Rather, CosMIC initially convolves the spike trains with a triangular pulse (**B**). The STC first discretizes the temporal interval and utilises the counts of spikes in each time bin, the bin edges and counts are plotted in **C**. The SR uses a bin centered around each true spike — an estimate in that bin is deemed a true detection (**D**). In order that the metric scores are comparable, we fix the STC and SR bin widths to be equal to CosMIC’s pulse width.

### 4.2 Spike train correlation

The first step in the calculation of the spike train correlation (STC) is the discretization of the temporal interval into bins of width *Δ*_2_. Two vectors of spike counts, **c** and **ĉ**, are subsequently produced, whose *i*^th^ elements equal the number of spikes in the *i*^th^ time bin for the true and estimated spike trains, respectively. The STC is the Pearson product-moment correlation coefficient of the resulting vectors, i.e.

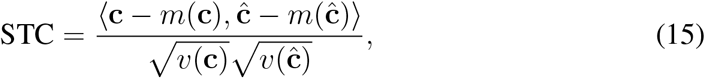

where 〈·, ·〉, *ν*(·) and *v*(·), represent the inner product, sample mean and sample variance, respectively. To remain consistent with the success rate, in all numerical experiments, we define *Δ*_2_ = *Δ*_1_ = *ϵ*.

The STC takes values in the range [−1, 1]. In practise, however, it is rare for a spike detection algorithm to produce an estimate that is negatively correlated with the ground truth (Berens et al., 2017). Moreover, an estimate with maximal negative correlation is equally as informative as one with maximal positive correlation. In this paper, we utilise the normalised spike train correlation, the absolute value of the STC. This ensures that the range of each metric that we analyse is equivalent (and equal to [0,1]) and that, as a consequence, the distribution of metric values are comparable.

### 4.3 Victor-Purpura dissimilarity

Victor and Purpura (1997) introduced a distance metric to compare the dissimilarity between sets of spikes from different neurons: 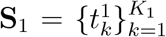 and 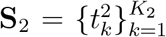 The distance is the minimum cost of transforming one set of spikes into the other using a set of three operations: insertion, deletion and temporal shifts of spikes. A cost is associated with each operation; insertion and deletion both carry a cost of 1, whereas the cost of a temporal shift depends on the extent of the shift and the value of a parameter, *q*. In particular, the cost of transforming one spike into another is

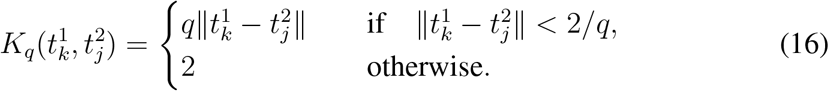

If the spikes are within the precision prescribed by the shift parameter, 2/*q*, the cost relates to a temporal shift. Otherwise, the cost invoked is the sum of the costs of deleting one spike and inserting another at the correct location. In all experiments, we set 2/*q* to be equal to CosMIC s pulse width, so that the minimum tolerated precision of CosMIC and this metric are equivalent. Finally, the distance between two sets of spikes, *D*_VP_ (S_1_, S_2_), is the minimum total cost of the operations transforming one spike train to the other. A larger score indicates less similar spike trains, whereas the minimum score, zero, is awarded to identical spike trains.

### 4.4 van Rossum dissimilarity

A distance metric introduced by van Rossum (2001) was also designed to quantify the dissimilarity between sets of spikes from different neurons. The respective spike trains are first convolved with a biologically-motivated pulse, *q*(*t*) = exp(−*t*/*τ*) 1_*t*>0_, where *τ* is a tunable parameter and 1 is the indicator function. The metric score is the Euclidean distance between the resulting pulse trains, *f*_1,*τ*_ and *f*_2,*τ*_, such that

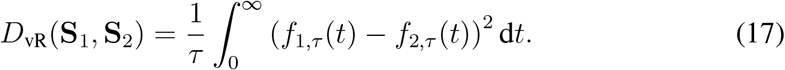

Following Kreuz et al. (2007), when computing the score of the van Rossum dissimilarity, we set *τ* with respect to the Victor-Purpura metric parameter: *τ* = 1/*q*.

## 5 Results

To investigate metric properties, we simulated estimated and ground truth spike trains and analysed the metric scores. To mimic the temporal error in spike time estimation, unless otherwise stated, estimates were normally distributed about the true spike times. In the following, we refer to the standard deviation of the normal distribution as the ‘jitter’ of the estimates.

### 5.1 CosMIC rewards high temporal precision

CosMIC was more sensitive to temporal precision than the STC or success rate (Fig. 5). First, we investigated this characteristic at the level of estimates of a single spike, *t*_true_. CosMIC depends only on the absolute difference between the estimate, *t*_est_, and the true spike — the further the distance, the smaller the score. The relationship between CosMIC and the temporal error, *Δ* = *t*_true_ − *t*_est_, is

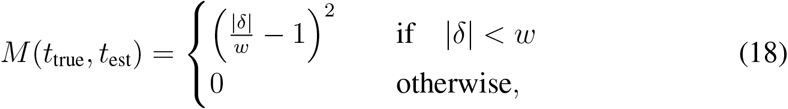

where *w* is the width of the pulse. The derivation of this result is given in Appendix A.2. The success rate, on the other hand, does not reward increasing temporal precision above the bin width; an estimate is assigned a score of 1 or 0, when its precision is above or below the bin width, respectively. Moreover, the STC is asymmetric in the temporal error; estimates the same distance from the true spike are not guaranteed to be awarded the same score, see Fig. 5A. This asymmetry stems from this metric’s temporal discretisation. The temporal interval is first discretised into time bins and the number of spikes in each bin are counted (Fig. 4). It follows that estimated spikes that are the same absolute distance from a true spike can fall into different time bins, thus achieving a different score. We note that the STC is always positive in Fig. 5A as, in this paper, we utilise the absolute value of the correlation (see Section 4.2).

**Figure 5:**
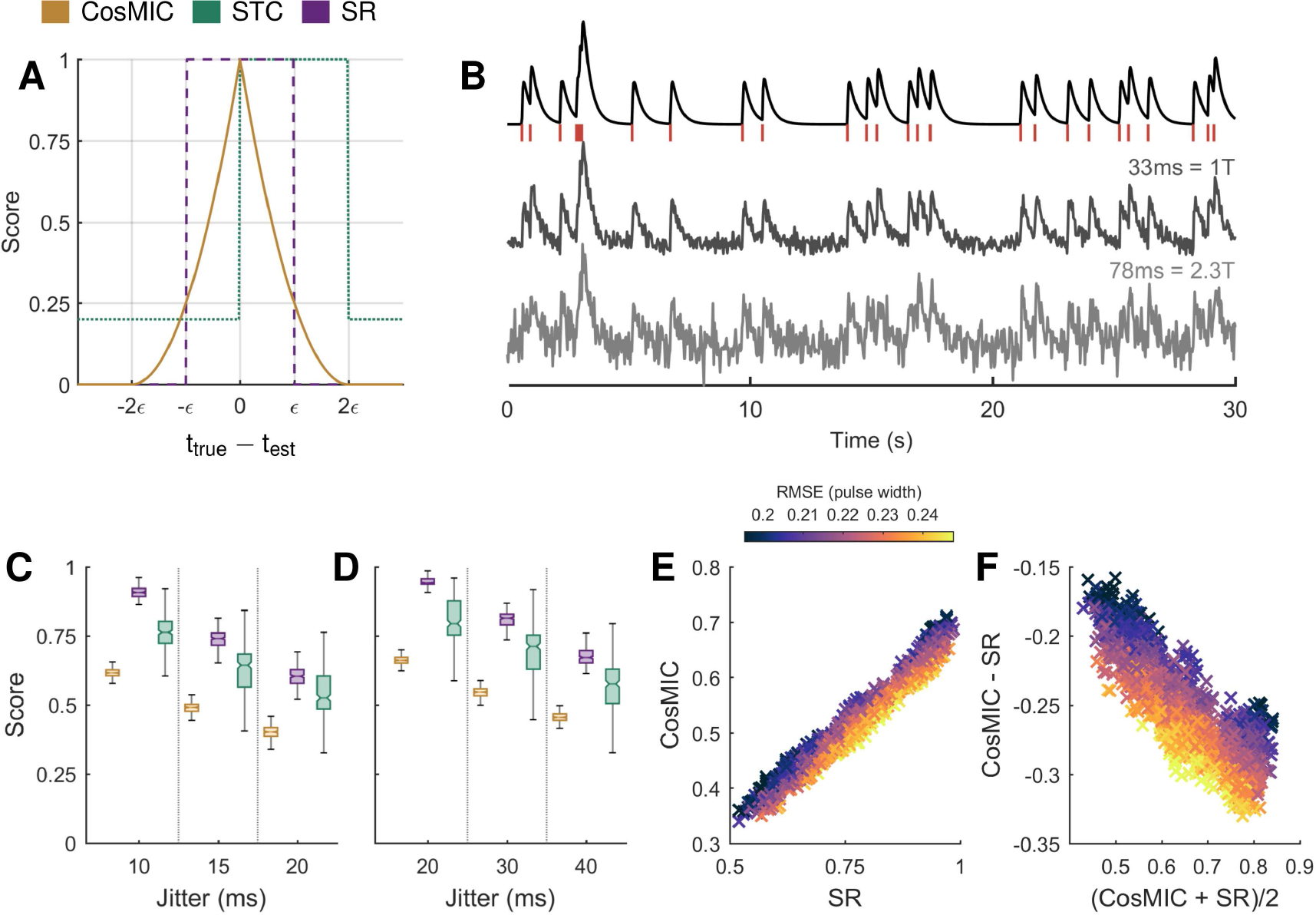
CosMIC was more sensitive to the temporal precision of estimates than the spike train correlation (STC) or success rate (SR). Unlike the STC, CosMIC awards estimated spikes (***t***_est_) with the same proximity to the true spike (***t***_true_) the same score (A). In contrast to both the STC and SR, CosMIC rewards increasing precision above the pulse width (**2**ϵ) with strictly increasing scores. In C and D, we plot the distribution of scores awarded to estimates that detect the correct number of spikes at varying temporal precision, in a low and high noise setting, respectively. In B, a sample of each of the following signals are plotted: the ground truth spike train, simulated as a Poisson process at rate 1Hz over 200s; the corresponding calcium transient signal, sampled with interval ***T*** =1/30s; the low and high noise fluorescence signal and the corresponding pulse widths. At each noise and jitter level, 100 realisations of spike train estimates normally distributed about the true spike times were generated. In both the low (C) and high noise (D) settings, the STC exhibited a relatively large variation in the scores awarded to estimates of the same jitter. CosMIC and the SR were roughly linearly related (E). CosMIC was boosted with respect to the success rate when temporal error, represented by the root mean square error (RMSE) of estimates as a fraction of the pulse width, was low (F). Conversely, CosMIC was relatively low with respect to the SR when temporal error was relatively high. The colormap in E and F is thresholded at the 1^st^ and 99^th^ percentiles of the RMSE for visual clarity.

On simulated data, we investigated the effect of these properties when spike train estimates, rather than single spikes, were evaluated. In particular, we analysed the metric scores when spike train estimates contained the correct number of spikes but their temporal precision varied. We simulated the ground truth spike train as a Poisson process with rate 1Hz over 200s. The corresponding calcium transient signal was generated assuming a Cal-520 pulse shape (see Table 1) and a sampling rate of 30 Hz. White Gaussian noise was added to the calcium transient signal to generate two fluorescence signals, one with low and the other with relatively high noise (Fig. 5B). The corresponding metric pulse widths, as calculated from the CRB, were 33ms and 78ms, or 1 and 2.3 sample widths, respectively. Spike train estimates were normally distributed about the true spikes with varying jitter. The metric scores were then calculated for 100 realisations of spike train estimates at each jitter level in both the low and high noise settings (Fig. 5C and D, respectively).

As the correct number of spikes were always estimated, the level of jitter represented the quality of a spike train estimate in this setting. Ideally, a metric would reliably reward spike train estimates of the same quality with the same score. The STC, however, took a relatively large range of values for estimates of the same jitter (Fig. 5C and D), despite having the same range as CosMIC and the success rate. This inconsistency is a consequence of the edge effects introduced by binning. Here, we use the term consistency in line with its semantic rather than mathematical definition.

We observed a roughly linear trend in the scores of CosMIC and the success rate (Fig. 5E). As expected, CosMIC was boosted with respect to the success rate when the root mean square error (RMSE) of detected spikes was relatively low when measured as a fraction of the pulse width. In each case, the RMSE was computed empirically from the estimated spikes within the precision of CosMIC and the success rate’s pulse width. Conversely, CosMIC was relatively low with respect to the success rate when the RMSE was relatively high. This trend is visible in the Bland-Altman plot (Altman and Bland, 1983; Giavarina, 2015), in which the mean of the two methods is plotted against the difference. We conclude that CosMIC is more sensitive to the temporal precision of detected spikes, as, unlike the success rate, it discriminates precision above the bin width.

### 5.2 CosMIC penalises overestimation

As opposed to the STC, CosMIC and the success rate penalised overestimation of spikes (Fig. 6). We simulated spike train estimates that were normally distributed about the true spike times. When the number of detected spikes (*K*_est_) was less than the number of true spikes (*K*_true_), the locations about which the estimates were distributed were chosen without replacement. When *K*_est_ > *K*_true_, the set of locations included all the true spikes plus a subset of extras chosen with replacement. The overestimation ratio (*K*_est_/*K*_true_) reflects the degree of accuracy to which an estimate matches the rate of a ground truth spike train. We observed that, rather than penalising overestimation, the STC increased with the overestimation ratio. In contrast, CosMIC and the success rate were maximised when the correct number of spikes were detected. This behaviour was consistent as the jitter of the estimated spikes varied; in this example, the jitter was *σ*_CRB_ (Fig. 6A), 2 *σ*_CRB_ (Fig. 6B) and 3 *σ*_CRB_ (Fig. 6C), respectively.

**Figure 6:**
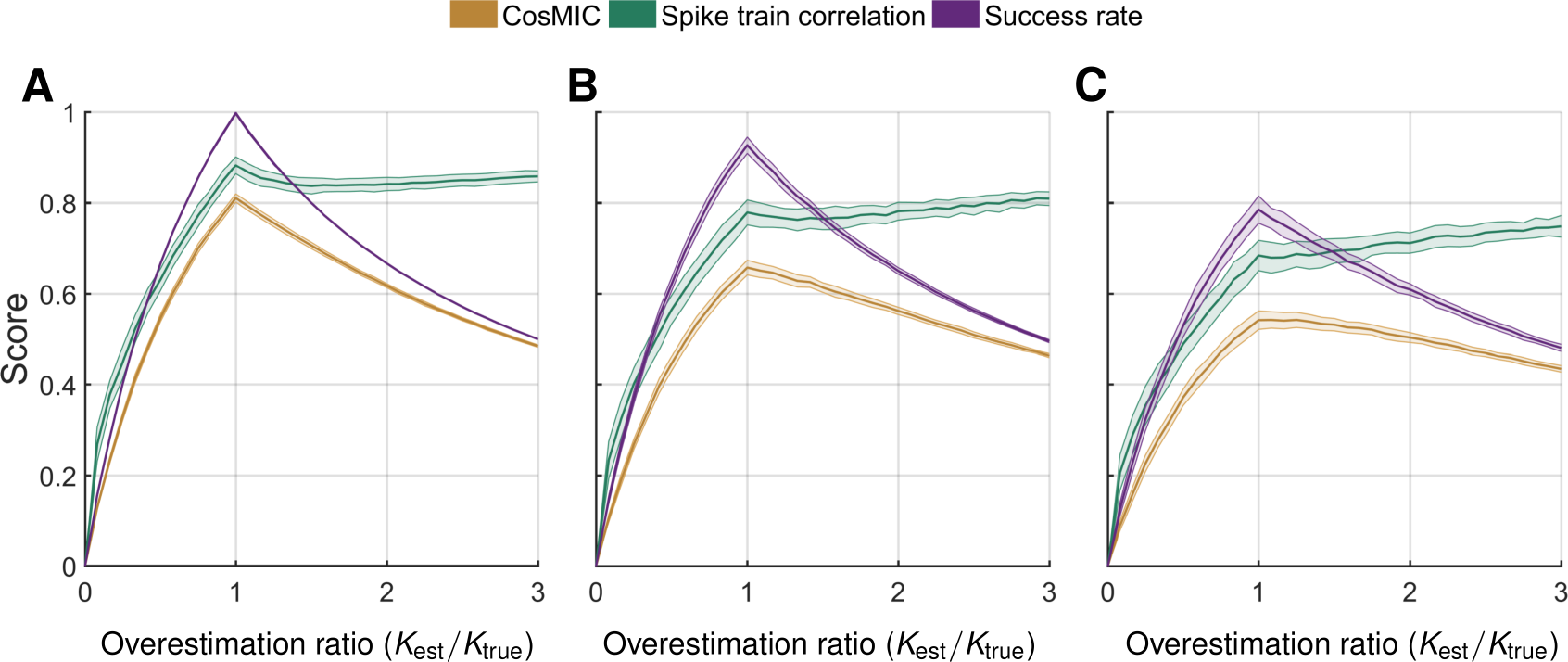
In contrast to the spike train correlation, CosMIC and the success rate were maximised when the correct number of spikes were detected. We display the distribution of metric’scores as the number of estimated spikes (*K*_est_) varies with respect to the number of true spikes (*K*_true_). The true spike train, which was identical throughout, consisted of 200 spikes simulated from a Poisson process with spike rate 1Hz. Estimated spikes were normally distributed about the true spikes, with jitter σ_CRB_ (A), 2 σ_CRB_ (B) and 3 σ_CRB_ (C), respectively, where σ_CRB_=20ms. When the number of estimated spikes was greater than the number of true spikes, estimates were distributed around a set of locations including all true spikes plus an extra subset chosen with replacement. For each metric we plot the mean (darker central line) and standard deviation (edges of shaded region) of metric scores on a set of 100 spike train estimates generated at each overestimation and jitter combination.

It is the type of normalisation used by the STC that caused it to be insensitive to overestimation. Scaling factors present in the spike count vectors cancel out in the numerator and denominator, see Eq. (15), rendering the STC invariant under scalar transformations of the inputs. When the STC was adapted to the continuous-time assessment of spike train similarity, by first convolving spike trains with a smoothing pulse, this flaw persisted (Paiva et al., 2010).

When the spike train estimates have jitter *σ*_CRB_ and their rate increases from perfect rate estimation to an overestimation ratio of 3, the success rate and CosMIC scores are reduced by 49% and 40%, respectively. Both metrics are thus penalising overestimation, with the former metric doing so more harshly. When the jitter is larger than the CRB, the reduction in CosMIC from perfect rate estimation to overestimation is relatively smaller, as CosMIC is already substantially penalising the temporal discrepancy.

### 5.3 Application to real imaging data

On imaging data of the mouse visual cortex at a frame rate of 13 Hz, CosMIC was more sensitive than the success rate to the temporal precision of detected spikes (Fig. 7). For a detailed description of the imaging data, see Reynolds et al. (2017). Briefly, four neocortical layer-5 pyramidal cells were simultaneously recorded in whole-cell configuration, different Poisson spiking patterns were evoked by brief current pulses, and calcium transients were imaged with a two-photon laser-scanning microscope (see Abrahams-son et al. 2017), thus establishing a realistic imaging data set with electrophysiological ground truth. An existing algorithm was used to detect spikes from each of 83 traces (Oñativia et al., 2013; Reynolds et al., 2016). Detected spike trains were subsequently compared to the electrophysiological ground truth using CosMIC, the success rate and the STC.

**Figure 7:**
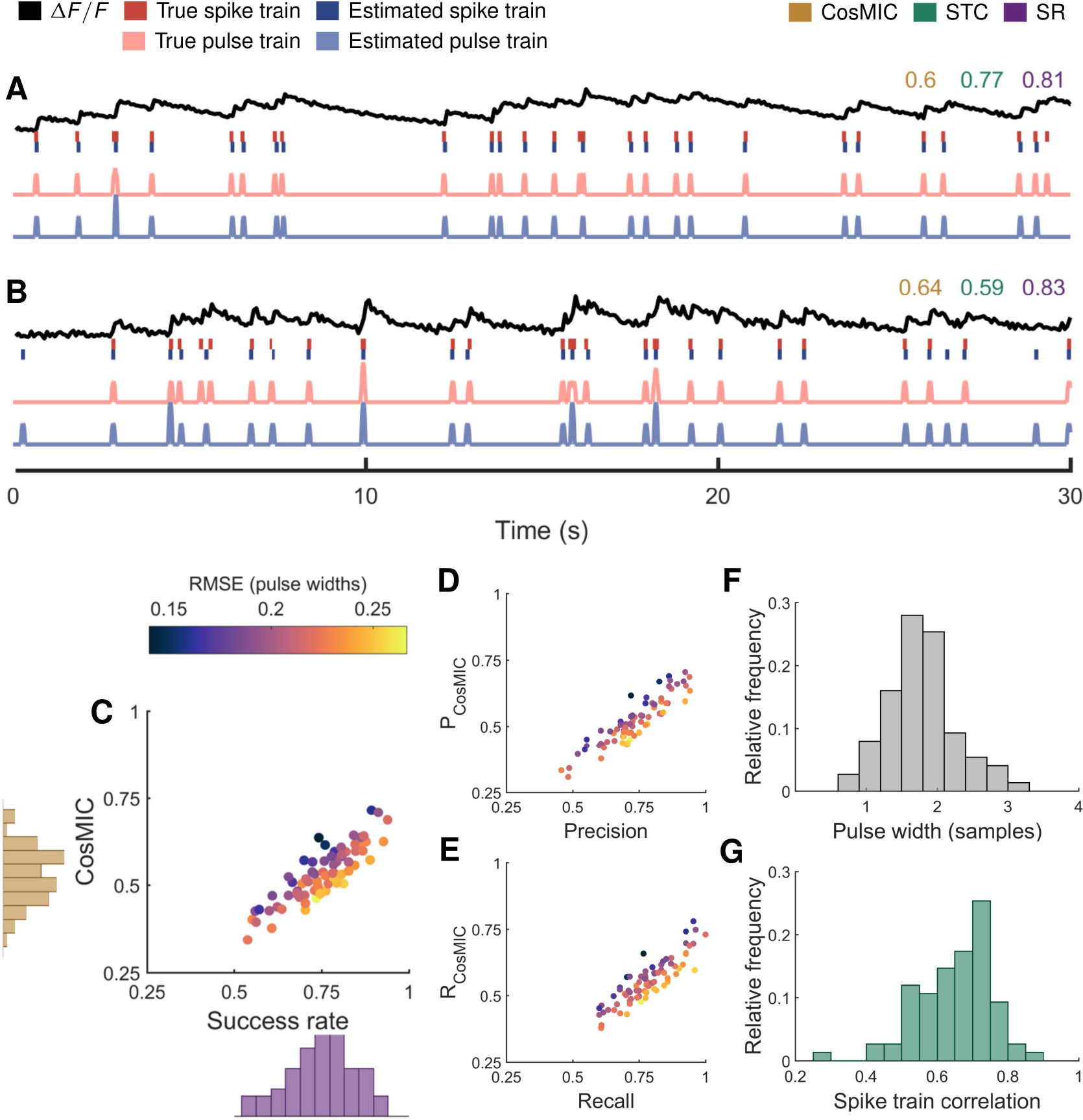
On mouse in vitro imaging data, CosMIC was more sensitive than the success rate (SR) to the temporal precision of detected spikes. Spikes were detected using an existing algorithm (Oñativia et al. 2013; Reynolds et al. 2016) from 83 traces sampled from visual cortex slices at 13Hz. In A and B, we display from top to bottom: an example fluorescence trace (*ΔF/F*), ground truth and detected spike trains, and the corresponding pulse trains. There was an approximately linear relationship between CosMIC and the SR (C). CosMIC was relatively high with respect to the SR when temporal error, represented by root mean square error (RMSE) as a fraction of the pulse width, was relatively low. Conversely, CosMIC was low with respect to the SR when temporal error was relatively high. This pattern was conserved in the relationship between the precision and CosMIC’s analogous ancestor metric, *P*_CosMIC_, (D) and between the recall and *R*_CosMIC_ (E). The range of pulse widths as computed from the Cramér-Rao bound (F) and the range of spike train correlation (STC) scores (G) are also shown.

As detailed in Section 3, the metric’s pulse width was set with respect to the CRB. On this dataset, the pulse widths were concentrated between 1 and 3 sample widths — this range encompassed 92% of the data, see (Fig. 7F). As the noise level of the data increases, so does the pulse width, see Eq. (11). Consequently, the tolerance of the metric with regards to the temporal precision of estimates also increases. As a result, estimates on noisier data (Fig. 7B) were scored with more lenience than those on less noisy data (Fig. 7A).

As was found on simulated data in Section 5.1, there was a linear trend between the scores of CosMIC and the success rate (Fig. 7C). CosMIC was relatively high with respect to the success rate when the temporal precision, represented by RMSE as a fraction of the pulse width, was relatively high. Conversely, CosMIC was low with respect to the success rate when the temporal precision was relatively low. This pattern was conserved when CosMIC’s ancestor metrics, *P*_CosMIC_ and *R*_CosMIC_ (Section 2.1), were compared to the precision and recall (Fig. 7D and E). The average RMSE over all traces was 27ms, or 0.37 sample widths. As CosMIC is able to discriminate precision above the pulse width, it is more able to reward this super-resolution performance than the success rate or STC.

### 5.4 CosMIC discriminates precision and recall of spike trains

By construction, CosMIC bears a strong resemblance to the Sørensen-Dice coefficient, which, in the context of spike detection, is referred to as the success rate. The success rate is the harmonic mean of the precision and recall, two intuitive metrics which represent the proportion of estimates that detect a ground truth spike and the proportion of true spikes detected, respectively. In this section, we demonstrate that CosMIC can accurately discriminate both the precision and recall of spike train estimates.

When a spike train estimate detects exactly a subset of the true spikes, plus no remainders, CosMIC and the success rate depend only on the percentage of true spikes detected (the recall) and not the location of that subset, see Fig. 8A and D. Denoting the size of the subset of true detections as *K* − *R*, with *K* the number of true spikes and 0 ≤ *R* ≤ *K*, we have

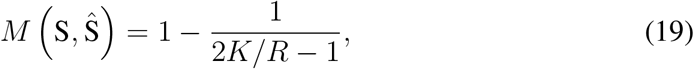

see Appendix A.4 for a proof. Thus, CosMIC depends only on the proportion of missing’ spikes, *R*/*K*, not their location. In contrast, the STC exhibited significant variation at each level of recall. This is illustrated in Fig. 8A, in which we plot the distribution of CosMIC, success rate and correlation scores over 100 realizations of spike train estimates at each level of recall. It can be seen that, in this setting, CosMIC and the success rate are fixed with the recall of the spike train estimates.

**Figure 8:**
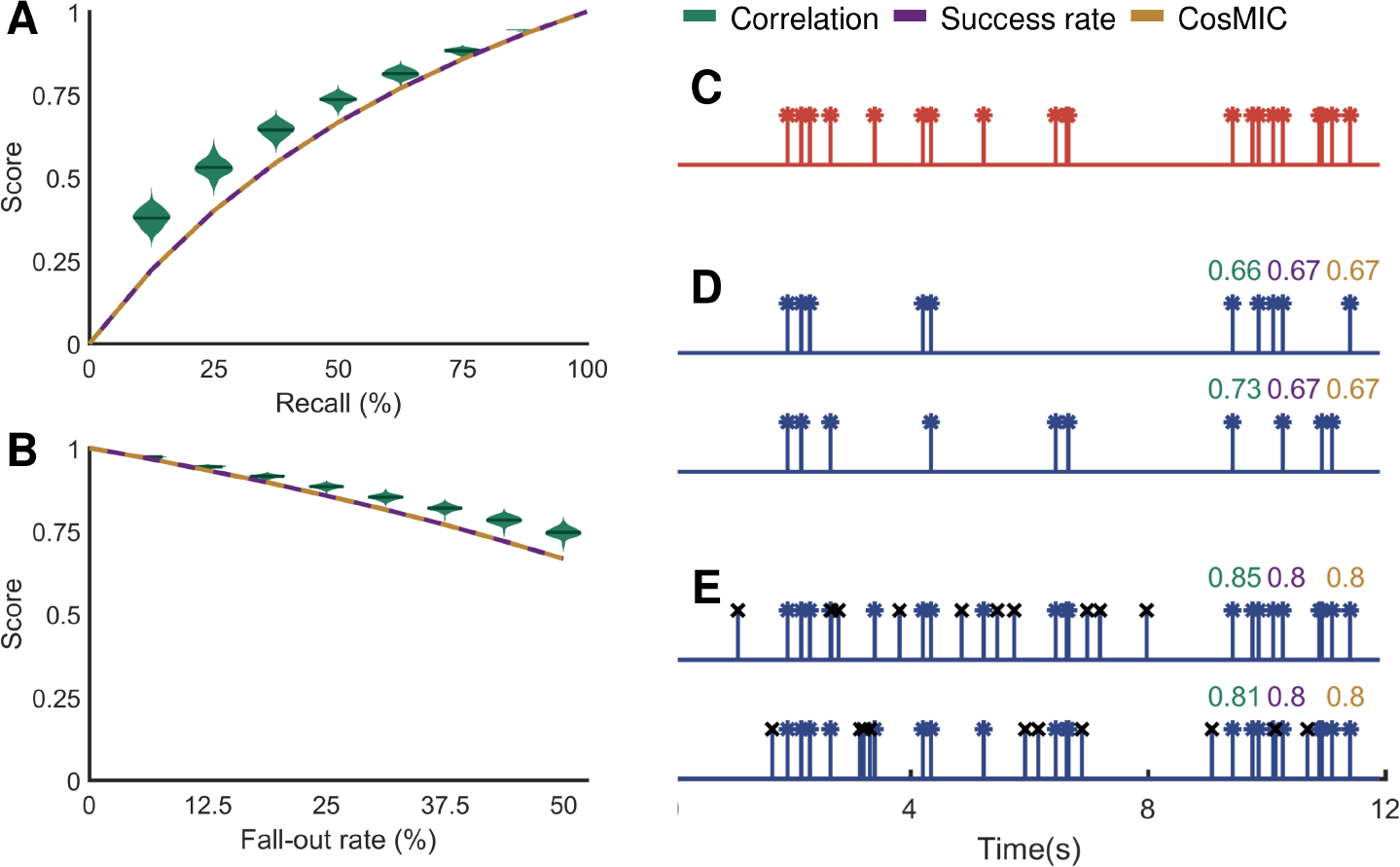
CosMIC’scored estimated spike trains of the same recall and fall-out rate consistently, unlike the spike train correlation (STC). When a spike train estimate detected precisely the location of a subset of spikes from a true spike train, the scores of CosMIC and the success rate depended only on the percentage of spikes detected (the recall), not the location of the detected spikes (**A, D**). In contrast, the STC varied with the subset of spikes that were detected. When a spike train estimate detected all the true spikes precisely plus a number of surplus spikes, the STC varied with the placement of the surplus spikes (**B, E**). In contrast, the success rate and CosMIC depended only on the percentage of estimated spikes that did not correspond to ground truth spikes (the fall-out rate, also known as the false positive rate). The distribution of correlation scores plotted in **D** and **E** stem from 100 realizations of estimated spike trains at each recall and fall-out rate. In **C**, we plot an example of a true spike train. In **D** and **E**, we plot estimated spike trains, with a recall and fall-out rate of 50% and 33%, respectively, along with the corresponding metric scores. The spikes with a black ‘x’ marker in **E** indicate the surplus spikes.

When all the true spikes were exactly detected plus *R* ≥ 0 surplus spikes, CosMIC and the success rate depend only on the level of precision not the location of the surplus spikes, see Fig. 8B and E. We have

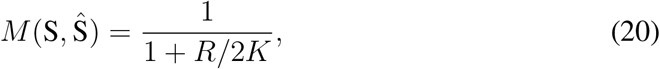

where *K* is the number of true spikes, see Appendix A.5 for a proof. The fall-out rate, which is the complement of the precision, is the proportion of estimates that were not deemed to have detected a ground truth spike. It is apparent from Eq. (20) that, in this setting, CosMIC depends only on the fall-out rate, *R*/*K*. The correlation, on the other hand, varied with the location of the surplus spikes. In Fig. 8E, we plot the distribution of the correlation scores for 100 realizations of spike train estimates at each level of precision. CosMIC and the success rate, which were constant (and identical) at a given precision, in this scenario, are also shown.

### 5.5 Comparison with Victor-Purpura and van Rossum distances

The Victor-Purpura (VP) and van Rossum (vR) spike distances were originally designed to quantify the dissimilarity between spike trains from different neurons (Victor and Purpura, 1997; van Rossum, 2001). Due to the obvious parallels between that scenario and ours, we investigated the applicability of the VP and vR metrics to scoring spike inference.

The vR metric initially convolves the respective spike trains with a causal exponential pulse and computes the Euclidean distance between the resulting pulse trains (Section 4.4). Despite the causality of the pulse, the metric’score is symmetric in the error of a single estimate about a true spike (Fig. 9A). The VP distance implicitly evokes a box function pulse, resulting in a piecewise linear relationship between the error of an estimate and the metric’score (Fig. 9A). Although the VP distance is not defined with respect to a smoothing pulse, this interpretation follows from an analogous argument to that presented in A.2. It is known that, as the pulse width increases from small to large with respect to the interspike interval, both metrics vary between coincidence detectors and rate detectors. To the best of our knowledge, the optimal pulse width for a compromise between rate and timing detection is not known, so we set the widths of vR and VP with respect to CosMIC’s pulse width.

**Figure 9:**
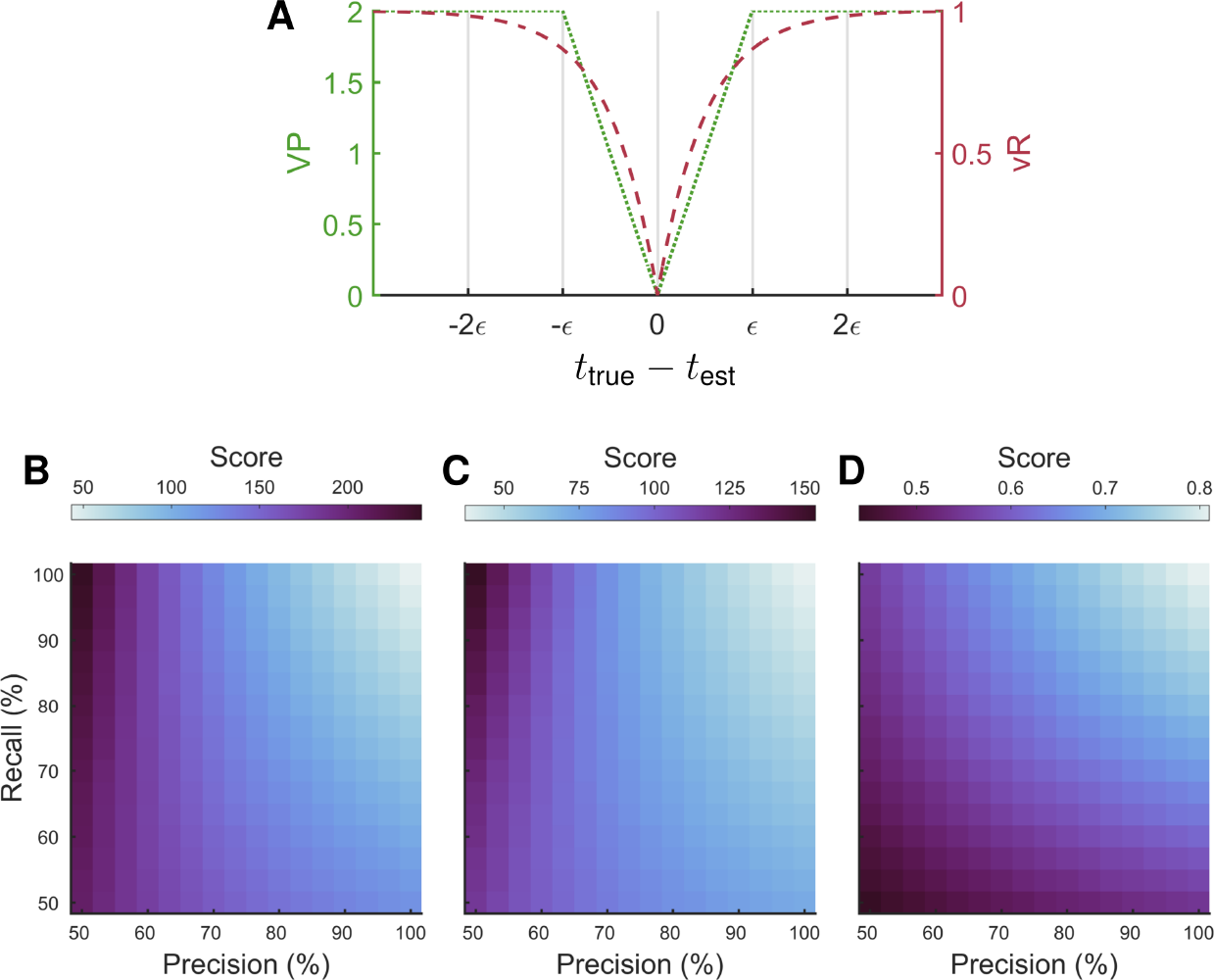
CosMIC was more sensitive to the precision and recall of spike train estimates than the Victor-Purpura (VP) or van Rossum (vR) spike distances. Both VP and vR are dissimilarity metrics, reaching a minimum of 0 when a true spike train and estimated spike train are equivalent. In A, this is demonstrated for one estimate (*t*_est_) of one spike (*t*_true_). The parameters of VP and vR were set with respect to CosMIC’s pulse width, **2**ϵ, which, in this example, was computed from a CRB of 20ms. The VP and vR distances were less sensitive to the recall than the precision of spike train estimates (B and C, respectively). CosMIC, however, only attained a relatively high score when both the precision and recall were high (D). At each level of precision and recall, the metric scores were averaged over 100 realisations of spike train estimates. The ground truth spike train contained 200 Poisson distributed spikes at rate 1Hz. False positives were uniformly distributed about the temporal interval, whereas true positives were normally distributed about true spikes with jitter 20ms.

Although it is already clear that, when the width is set correctly, VP and vR can discriminate the rate and temporal precision of spike trains with respect to one another (Paiva et al., 2010), it is not clear whether they are suitable for scoring spike train estimates. In Fig. 9B-D, we plot the scores of VP, vR and CosMIC, respectively, as the precision and recall of spike train estimates vary. We observed that vR and VP were less sensitive to the recall than the precision of spike train estimates; relatively low distances were obtained when only 50% of true spikes were detected. In contrast, CosMIC only attained a relatively high score when both the precision and recall were high (**D**). As it is crucial that a spike inference metric penalises both undetected and falsely-detected spikes, this result suggests that, without modification, VP and vR are not ideal for scoring spike train estimates.

The results correspond to a ground truth spike train consisting of 200 spikes generated from a Poisson process with rate 1Hz. False positives were uniformly distributed about the temporal interval, whereas true positives were normally distributed about true spikes with jitter 20ms. The pulse width was set assuming a CRB of 20ms. At each level of precision and recall, results were averaged over 100 realisations of spike train estimates.

## 6 Discussion

Much recent attention has been focused on the development of algorithms to detect spikes from calcium imaging data, while the suitability of the metrics that assess those algorithms have been predominantly overlooked. In this paper, we presented a novel metric (‘CosMIC’) to assess the similarity of spike train estimates compared to the ground truth. Our results demonstrate that CosMIC accurately discriminates both the temporal and rate precision of estimates with respect to the ground truth.

Using two-photon calcium imaging, the activity of neuronal populations can be monitored in vivo in behaving animals. Inferred spike trains can be used to investigate neural coding hypotheses, by analysing the rate and synchrony of neuronal activity with respect to behavioural variables. To justify such analysis, the ability of spike detection algorithms to generate accurate spike train estimates must be verified. When spike frequency is to be investigated, it is crucial that an estimate accurately matches the rate of the ground truth spike train. We have shown that the STC is not fit for this purpose; rather than penalising overestimation of the number of spikes, it is rewarded (Fig. 6). In contrast, CosMIC and the success rate are maximised when the correct number of spikes are detected. When the ultimate goal is to analyse spike timing with respect to other variables, it is critical that spikes can be detected with high temporal precision. We have shown that CosMIC has superior discriminative ability in this regard, compared to the success rate and STC (Fig. 5).

The current inconsistency in the metrics used to assess spike detection algorithms, hinders both experimentalists, aiming to select an algorithm for data analysis, and developers. In light of this problem, a recent benchmarking study tested a range of algorithms on a wide array of imaging data (Berens et al., 2017). Although informative, the study, which relied heavily on the STC to assess algorithm performance, may not provide the full picture. By introducing a new metric, we hope to complement such efforts in the pursuit of a thorough, quantitative evaluation of spike inference algorithms.

By construction, CosMIC bears a resemblance to the Sørensen-Dice coefficient, which is commonly used to compare discrete, presence/absence data (Dice, 1945; Sørensen, 1948). This metric, which is also known as the F1-score, is widely used in many fields, including ecology (Bray and Curtis, 1957) and image segmentation (Zou et al., 2004). When applied to spike inference, this coefficient is referred to as the success rate and is one of the two most commonly used metrics. We have demonstrated that this construction confers some of the advantages of the success rate to CosMIC. In particular, CosMIC is able to accurately discriminate the precision and recall of estimated spike trains (Fig. 8). We have also shown the advantages of CosMIC over the success rate; most importantly, it is more sensitive to a spike train estimate’s temporal precision than the success rate (Fig. 5). Furthermore, CosMIC’s parameter is defined with respect to the statistics of the dataset and unlike the success rate’s bin size, it does not need to be selected by a user.

We demonstrated that CosMIC is boosted with respect to the success rate when temporal precision is relatively high. In particular, as temporal precision approaches the CRB, CosMIC increases to a maximum. It is not clear how close existing algorithms are to this theoretical bound. Nevertheless, it is important to discriminate between the temporal precision of algorithms, even if the performance is not yet optimal. For example, if all algorithms produce estimates with error on the order of a sample width, it is still of interest to know which algorithm produces the lowest error. With its graded pulse shape, CosMIC is able to penalise decreasing error in this way.

The width of the pulse is computed from a lower bound on temporal precision (Section 3), which, in turn, is derived from the statistics of the dataset. As a result, the metric will be more lenient for spike inference algorithms on noisier or lower sampling rate data. This is due to our assumption that a metric score should reflect the difficulty of the spike inference problem. To calculate the bound, knowledge of the calcium transient pulse parameters and the standard deviation of the noise are required. These parameters are typically used by algorithms in the spike detection process (Vogelstein et al., 2010; Deneux et al., 2016). Using only one pulse amplitude parameter, which relates to the amplitude of a single spike, is a simplification. Depending on the fluorescent indicator, amplitudes do, in fact, decrease (Liitcke et al., 2013) or increase (Chen et al., 2013) at high firing rates. Consequently, CosMIC may be slightly more punitive in the former case than the latter.

The problem of comparing a ground truth and estimated spike train is analogous to that of comparing spike trains from different neurons. In the spike train metric literature, binless measures have been found to outperform their discrete counterparts (Paiva et al., 2010). It is also common to convolve spike trains with a smoothing pulse prior to analysis (van Rossum, 2001; Schreiber et al., 2003). In that context, the width and shape of the pulse reflect hypotheses about the relationship between neuronal spike trains. A width that is large with respect to the average interspike interval results in a metric tuned to the comparison of neuronal firing rates. Conversely, a relatively small width produces a metric that acts as a coincidence detector. To apply CosMIC to the problem of spike train comparison, one could similarly vary the pulse width to tailor its performance to the neural coding scheme. In the context of spike detection, which we view as a parameter estimation problem, the pulse width is fixed with respect to a lower bound on the precision with which a spike time can be estimated. Setting the width via this bound, which is tailored to calcium imaging data, results in a metric that assesses how accurately parameters have been estimated given the constraints of the data. This approach would need to be altered to extend CosMIC to other applications. We note that, in the absence of this pulse width, CosMIC is sufficiently universal to be applied to the comparison of any point processes.

Finally, we note that the developed metric is able to accurately assess an estimate s temporal and rate precision. This information is unified in a single score that summarises the overall performance of an algorithm. We consider a single summary score to be practical for users who do not have the time or desire to analyse multi-dimensional trade-offs. Alternatively, CosMIC’s ancestor metrics, *R*_CosMIC_ and *P*_CosMIC_, can be used to determine the extent to which errors stem from undetected or falsely-detected spikes.

## Acknowledgments

This work was supported by European Research Council starting investigator award [grant number 277800] (Pier Luigi Dragotti); Biotechnology and Biological Sciences Research Council [grant number BB/K001817/1] (Simon R. Schultz); EU Marie Curie FP7 Initial Training Network [grant number 289146] (Simon R. Schultz); CIHR New Investigator Award [grant number 288936] (P. Jesper Sjöström); CFI Leaders Opportunity Fund [grant number 28331] (P. Jesper Sjöström); CIHR Operating Grant [grant number 126137] (P. Jesper Sjöström) and NSERC Discovery Grant [grant number 4185462] (P. Jesper Sjöström).

## A Appendices

In the appendices, we provide derivations of some results presented in the main text. The following notation is consistent throughout. We denote with *x*(*t*) and *X̂*(*t*) the true and estimated spike trains, see Eq. (1). We denote the triangular smoothing pulse with *p*_*ϵ*_(*t*), see Eq. (2). The true and estimated pulse trains are denoted *y*(*t*) = *x*(*t*) ∗ *p*_*ϵ*_(*t*) and *ŷ*(*t*) = *X̂*(*t*) ∗ *p*_*ϵ*_(*t*), respectively. The proposed metric score, when comparing the similarity between a ground truth set of spikes, 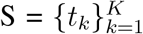, with a set of estimates, 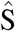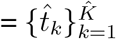, is

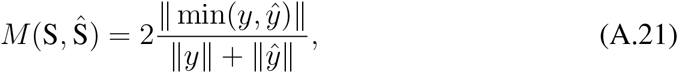

where ‖ · ‖ is the L1-norm.

## A.1 Alternative metric form

In the following, we derive an alternative equation for CosMIC; we show that

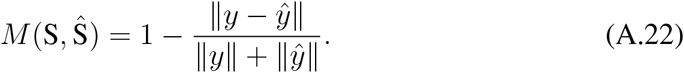

We have

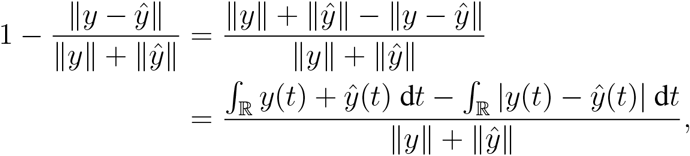

where we have used the fact that *y*(*t*) and *ŷ*(*t*) are non-negative for all *t* ∊ ℝ. Decomposing both integrals over ℝ into their counterparts over the disjoint sets {*t* ∊ ℝ: *y*(*t*) > *ŷ*(*t*)} and {*t* ∊ ℝ: *y*(*t*) ≤ *ŷ*/(*t*)} and subsequently combining them, we have

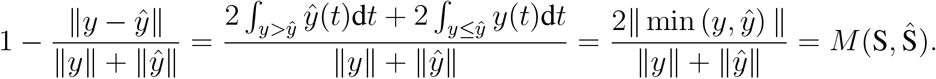

Eq. (A.22) then follows.

## A.2 Score for estimate of one spike

We now derive an expression for the metric score of the estimate of the location of one spike in terms of the temporal error of the estimate, |*u*|. We see that, as the temporal precision increases above the threshold precision (*ϵ*), the metric score increases monotonically.

**Proposition 1.** *The score given to an estimate of the location of a single spike, t*_0_, *with temporal error u* ∊ ℝ *is*

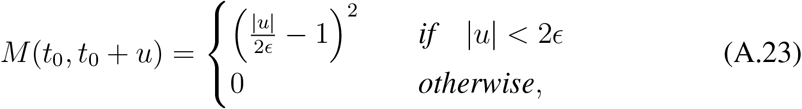

*where ϵ is half the width of the pulse, p*_*ϵ*_(*t*), *as in* *Eq.* (2).

*Proof.* Without loss of generality, we let the true spike location be at *t*_0_ = 0, as the metric score depends on the relative rather than absolute locations of the estimated and ground truth spikes. From Eq. (A.21), we have

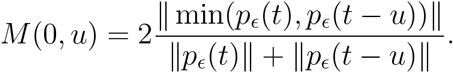

When |*u*| > 2*ϵ*, the pulses do not overlap and, consequently, the numerator is equal to 0. Therefore, the metric score is zero for all |*u*| > 2*ϵ*. For |*u*| < 2*ϵ*, we write

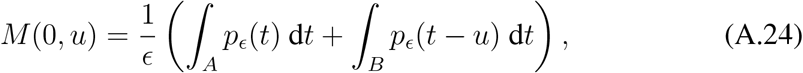

which follows from ‖*p*_*ϵ*_‖ = *ϵ*, *A* = {*t* ∊ ℝ: *p*_*ϵ*_(*t*) < *p*_*ϵ*_(*t* – *u*)} and *B* = {*t* ∊ ℝ: *P*_*ϵ*_(*t*) ≥ *p*_*ϵ*_(*t* – *u*)}. From the change of variables *v* = *t* + *u*, we see that *M*(0, *u*) = *M*(0, −*u*). As *M* is even in the second argument, we must only calculate *M*(0, *u*) for 0 < *u* < 2*ϵ*. To identify the support of *A* and *B*, we must identify the point at which *P*_*ϵ*_(*t*) = *P*_*ϵ*_(*t* – *u*). We have

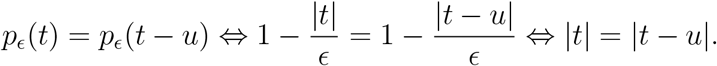

For 0 < *u* < 2*ϵ*, the intersection point occurs in the right half of *p*_*ϵ*_(*t*) and the left half of *p*_*ϵ*_(*t* – *u*), it follows that *t* = *u*/2. Eq. (A.24) becomes

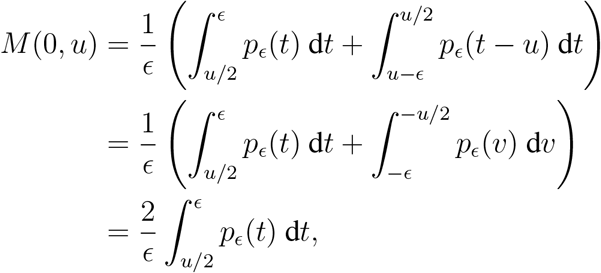

which follows from the change of variables *v* = *t* + *u* and the symmetry of *p*_*ϵ*_(*t*) about 0. Evaluating the integral, we obtain *M*(0, *u*) = (|*u*|/2*ϵ* – 1)^2^, for |*u*| < 2*ϵ*.

## A.3 Metric score at precision of CRB

The CRB is commonly used as a benchmark for algorithm performance in parameter estimation problems. In the context of calcium imaging, it has been previously used to evaluate detectability of spikes under different imaging modalities (Reynolds et al., 2015; Schuck et al., 2018). In this case, the CRB reports the minimum uncertainty achievable by any unbiased estimator when estimating the location of one spike. We thus set the width of the pulse to ensure that, on average, an estimate of the location of one spike at the precision of the CRB achieves a metric score of 0.8. This benchmark score is relatively high in the range of the metric, which is between 0 and 1, whilst allowing leeway to be exceeded.

**Proposition 2.** *Let t*_0_ *denote the location of the true spike. The estimate is normally distributed about the true spike at the precision of the CRB, it is modelled with the random variable* 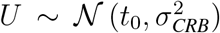. *We denote β* = *σ*_CRB_/*w*, *where w is the pulse width. Then, we have* 𝔼[*M*(*t*_0_, *U*)] = 0.8 *if β satisfies the following equation*,

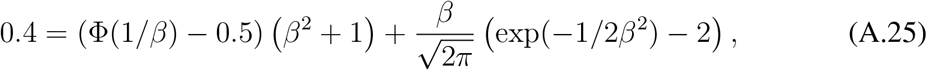

*where* Φ *denotes the cumulative distribution function of the standard normal distribution*.

*Proof.* We want to identify the pulse width at which 0.8 = 𝔼[*M*(*t*_0_, *U*)]. Without loss of generality, we consider the case where *t*_0_ = 0. Due to the fact that *M*(0, ·) is even and the results of Appendix A.2, we have

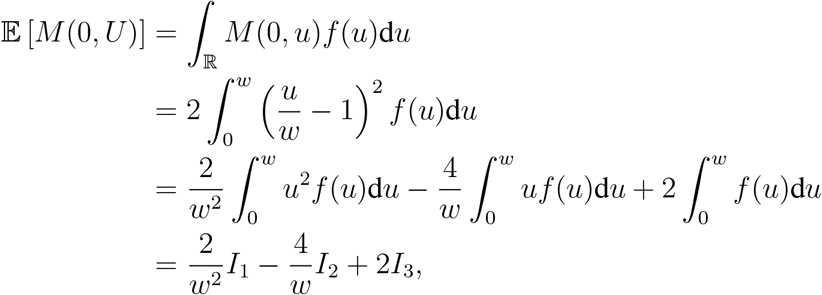

where *f*(·) is the probability density function of *U*. Applying integration by parts to *I*_1_, we obtain

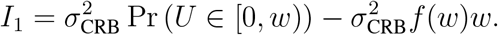

The remaining integrals are 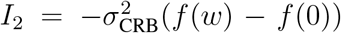 and *I*_3_ = Pr(*U* ∊ [0, *w*)), respectively. Putting the integrals together:

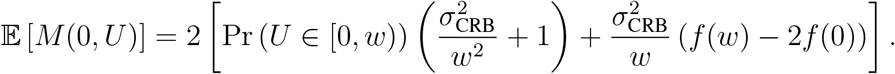

Writing *β* = *σ*_CRB_/*w*, we have

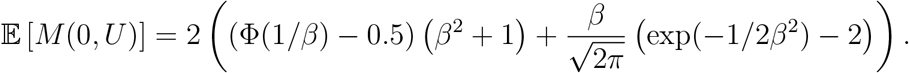

## A.4 Exact detection of subset of true spikes

We have a set of *K* true spikes, *S*, and *K̂* estimates, *Ŝ*. The set of estimates contains a subset of the ground truth spike times with the exception of *R* missing spikes and no extras, such that *K* = *K* – *R* with 0 ≤ *R* ≤ *K*. Due to the distributivity of the convolution operation

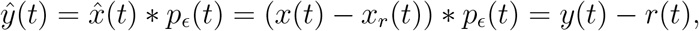

where *x*_*r*_(*t*) and *r*(*t*) are the spike train and pulse train, respectively, of the spikes missing from *Ŝ*. From the form in Eq. (A.22), the metric score becomes

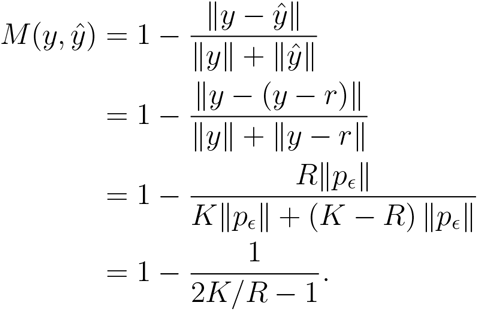

## A.5 Exact detection of all true spikes with overestimation

We have a set of *K* true spikes, S, and *K̂* estimates, *Ŝ*. The set of estimates contains all the ground truth spike times plus *R* ≥ 0 extra spikes, such that *K̂* = *K* + *R*. Due to the distributivity of the convolution operator, the estimated pulse train can be written

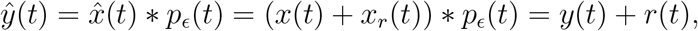

where *x*_*r*_(*t*) and *r*(*t*) are the spike train and pulse train, respectively, of the surplus spikes. From the form in Eq. (A.22), the metric score becomes

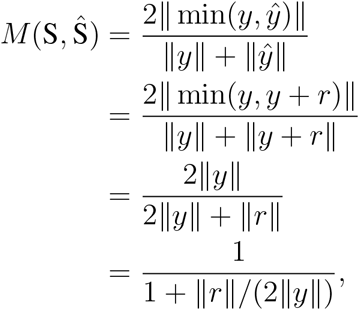

where the penultimate line follows from the non-negativity of *y* and *r*. As ‖*y*‖ = *K*‖*p*_*ϵ*_‖ and ‖*r*‖ = *R*‖*p*_*ϵ*_‖, it follows that

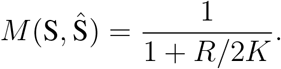

